# Bacteria conjugate ubiquitin-like proteins to interfere with phage assembly

**DOI:** 10.1101/2023.09.04.556158

**Authors:** Jens Hör, Sharon G. Wolf, Rotem Sorek

## Abstract

Multiple immune pathways in humans conjugate ubiquitin-like proteins to virus and host molecules as a means of antiviral defense. Here we studied an anti-phage defense system in bacteria, comprising a ubiquitin-like protein, ubiquitin-conjugating enzymes E1 and E2, and a deubiquitinase. We show that during phage infection, this system specifically conjugates the ubiquitin-like protein to the phage central tail fiber, a protein at the tip of the tail that is essential for tail assembly as well as for recognition of the target host receptor. Following infection, cells encoding this defense system release a mixture of partially assembled, tailless phage particles, and fully assembled phages in which the central tail fiber is obstructed by the covalently attached ubiquitin-like protein. These phages exhibit severely impaired infectivity, explaining how the defense system protects the bacterial population from the spread of phage infection. Our findings demonstrate that conjugation of ubiquitin-like proteins is an antiviral strategy conserved across the tree of life.

## Introduction

Ubiquitin is a conserved eukaryotic protein that can be covalently attached to target proteins as part of the protein degradation pathway^1^. Covalent attachment of ubiquitin necessitates a cascade of enzymatic reactions carried out by three classes of proteins called E1, E2 and E3^1^. The E1 enzyme first adenylates a conserved C-terminal glycine residue in the ubiquitin protein and then covalently attaches the adenylated ubiquitin to a conserved cysteine in the E1 active site^1^. The ubiquitin is then transferred to a cysteine residue in the E2 enzyme, which further transfers it to the target protein, usually via a mediator E3 protein^1^. Ubiquitination can be reversed by deubiquitinases (DUBs), which are peptidases capable of removing ubiquitin from target molecules^2^.

While ubiquitination of proteins is central to the protein degradation pathway in humans^3^, conjugation of ubiquitin and ubiquitin-like proteins (Ubls) was also shown to be important in pathways of innate immunity^4,5^. For example, interferon-stimulated gene 15 (ISG15) is a Ubl comprised of two fused ubiquitin-like domains, which is involved in protecting human cells against a variety of viruses such as influenza A and HIV-1^6^. ISG15 is one of the most highly upregulated genes following virus-induced type I interferon stimulation^7^, and its specific E1, E2, and E3 enzymes are also interferon-induced^8^. It was shown that upon virus infection, ISG15 is intracellularly conjugated to multiple cellular and viral protein targets, impairing viral propagation via a mechanism that is still unclear^6,9,10^.

Like eukaryotes, bacteria can also be infected by viruses (phages), against which bacteria have evolved a plethora of immune defense strategies^11^. A recent screen for anti-phage defense systems in bacteria revealed a four-gene operon encoding a Ubl protein, a protein with a predicted E1 domain, a protein with a predicted E2 domain, and a protein with a predicted DUB domain^12^ (Figure 1A). Similar to ISG15, the Ubl in this bacterial system is comprised of two fused ubiquitin domains, and the operon was accordingly denoted BilABCD (standing for <u>b</u>acterial ISG15-like system). The Bil defense system was previously shown to protect bacteria against multiple phages, but the mechanism of defense remained unknown^12^.

**Figure 1.**
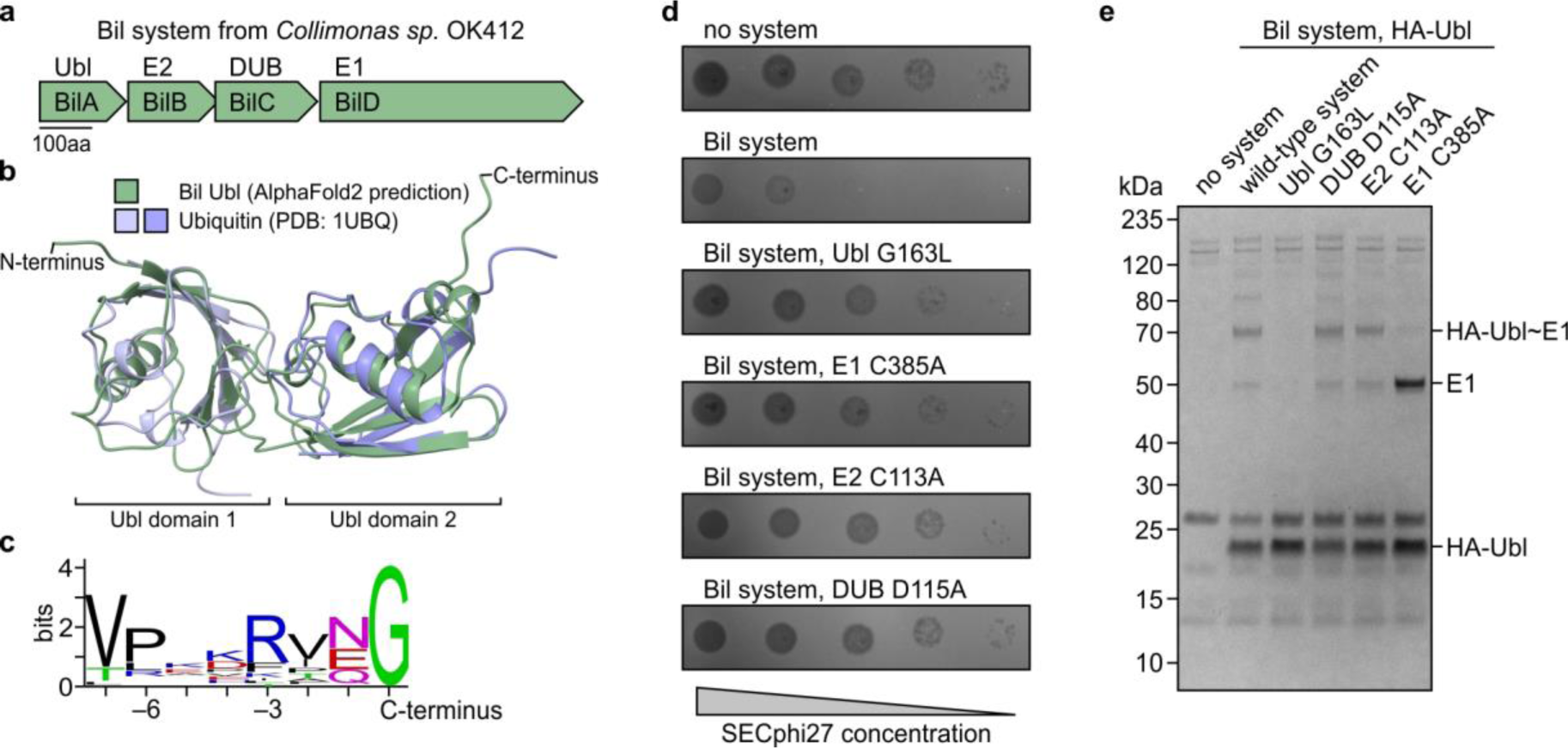
Features of ubiquitin-like conjugation in the Bil anti-phage defense system. **a,** Operon structure of the Bil system from *Collimonas sp.* OK412. **b,** Structure of the BilA Ubl protein, predicted by AlphaFold2^15^. The N- and C-terminal Ubl domains were each aligned separately to human ubiquitin (PDB: 1UBQ^16^) with RMSD values of 1.62 Å and 1.77 Å, respectively. **c,** C-terminal sequence conservation among homologs of the Ubl protein of the Bil system. **d,** Plaque assays showing the defense phenotype of the wild-type and mutant Bil systems against phage SECphi27. Data are representative of three independent experiments. **e,** Immunoprecipitation of HA-tagged Ubl protein from wild-type and mutant Bil system-expressing bacteria, analyzed using SDS-PAGE followed by Coomassie staining.

In this study, we show that the Bil system covalently conjugates its Ubl protein to the phage central tail fiber, an essential structural component of the phage tail tip that is crucial for both tail assembly^13^ and host recognition^14^. Conjugation of the Ubl to the central tail fiber inhibits tail assembly, leading to the production of non-infective, tailless phages. In some cases, phages manage to assemble fully tailed particles, but in these particles the Ubl is covalently attached to the central tail fiber at the tip of the tail, impairing infectivity likely by inhibiting host recognition. Thus, the Bil system prevents the spread of phages to neighboring cells after the initially infected bacterium is lysed, and protects the bacterial population as a whole. Our study reveals a new mechanism used by bacteria to defend against phage predation and sheds light on the mechanism of Ubl-mediated immunity.

## Results

### Functional features of Ubl conjugation in a bacterial defense system

We set out to study the BilABCD system from *Collimonas sp.* OK412, a four-gene operon encoding the Ubl protein BilA, the E2 domain protein BilB, the DUB BilC and the E1 domain protein BilD (Figure 1A). When heterologously expressed in *Escherichia coli*, this system was previously shown to protect against propagation of multiple coliphages^12^. Structural analysis using AlphaFold2^15^ revealed that the bacterial Ubl protein contains two fused ubiquitin-like domains, both of which are structurally highly similar to human ubiquitin (Figure 1B). The last amino acid at the C-terminus of the bacterial Ubl is glycine, which is fully conserved among all homologs of the BilA Ubl (Figure 1C). As the C-terminal residue of Ubls is an obligatory glycine whose carboxyl group is the site of conjugation to targets^1^, conservation of glycine at the C-terminus of the bacterial Ubl suggests that, similar to other known Ubl systems, conjugation to the target occurs via this residue. Indeed, a single amino acid substitution at this position (G163L) was sufficient to abolish phage defense by the Bil system (Figure 1D).

The first step in any Ubl conjugation cascade involves adenylation of the Ubl, followed by formation of a covalent thioester bond between the E1 enzyme and the Ubl^17^. To examine whether a covalent Ubl∼E1 complex is formed by the bacterial system, we tagged the E1 protein in the system and performed western blot analysis on total protein extracted from cells expressing the Bil system (Figure S1A). This analysis revealed two bands for the E1 protein, one of the expected size of free tagged E1 (∼53 kDa) and one that was ∼20 kDa larger than the free E1, suggesting linkage to a single Ubl (∼18 kDa). The addition of dithiothreitol (DTT), which reduces thioester bonds^18^, resulted in the disappearance of this additional band, further supporting the hypothesis that this band represents a Ubl∼E1 covalent complex (Figure S1A).

Immunoprecipitation of an N-terminally HA-tagged Ubl further confirmed that a Ubl∼E1 complex is formed in cells expressing the Bil system (Figure 1E). This covalent Ubl∼E1 complex was lost when the conserved cysteine at the active site of the E1 protein (C385) was substituted for alanine (Figure 1E). The C385A substitution also abolished the ability of the Bil system to protect against phages^12^, indicating that conjugation of the Ubl by the E1 enzyme is essential for the defensive function (Figure 1D, Table S6). Structural analysis with AlphaFold-Multimer^19^ predicted a high-confidence interaction between the two proteins that involves the penetration of the Ubl C-terminus into the E1 adenylation site (Figure S1B). This places the C-terminal glycine of the Ubl at the nucleotide-binding loop of the E1, thus providing a structural explanation for the mechanism of the first step of the Ubl∼E1 conjugation (Figure S1B). The structure further predicts a substantial non-covalent interaction interface between the E1 and the Ubl, explaining why the Ubl can pull down the C385A E1 protein even in the absence of covalent interactions (Figure 1E).

The second step in the Ubl conjugation cascade is the transfer of the Ubl from the cysteine of the E1 to a cysteine of the E2^17^. We were unable to detect a thioester-linked Ubl∼E2 complex, suggesting that this complex might be transient (Figure S1C). However, amino acid substitution in the predicted active site of the E2 (C113A) abrogated defense against phages, as previously shown^12^, further supporting the hypothesis that Ubl conjugation is central for the defensive function of the Bil system (Figure 1D). Disruption of the predicted active site of the DUB (D115A) also abrogated phage defense of the Bil system, suggesting that its deubiquitination enzymatic activity is essential for the function of the system (Figure 1D).

### The Bil system interferes with phage assembly

We next conducted liquid culture infection assays with SECphi27, a phage from the *Drexlerviridae* family against which the Bil system provides substantial protection in plaque assay experiments (Figure 1D). Cells expressing the Bil system survived phage infection in liquid culture when phages were supplied at low multiplicity of infection (MOI), but when infected at high MOI the culture completely collapsed (Figure 2A). These culture dynamics are similar to those observed for abortive infection (Abi) defense systems, which function by inducing death of infected cells at a time point earlier than that necessary for completion of the phage replication cycle^20^. However, cells expressing the Bil system that were infected at high MOI lysed at the same time as control cells rather than lysing at an earlier time point as expected for Abi systems^20^ (Figure 2A). This observation pointed to the possibility that the Bil system allows the phages to complete their temporally controlled replication cycle, but still somehow prevents the production of infective phage progeny.

**Figure 2.**
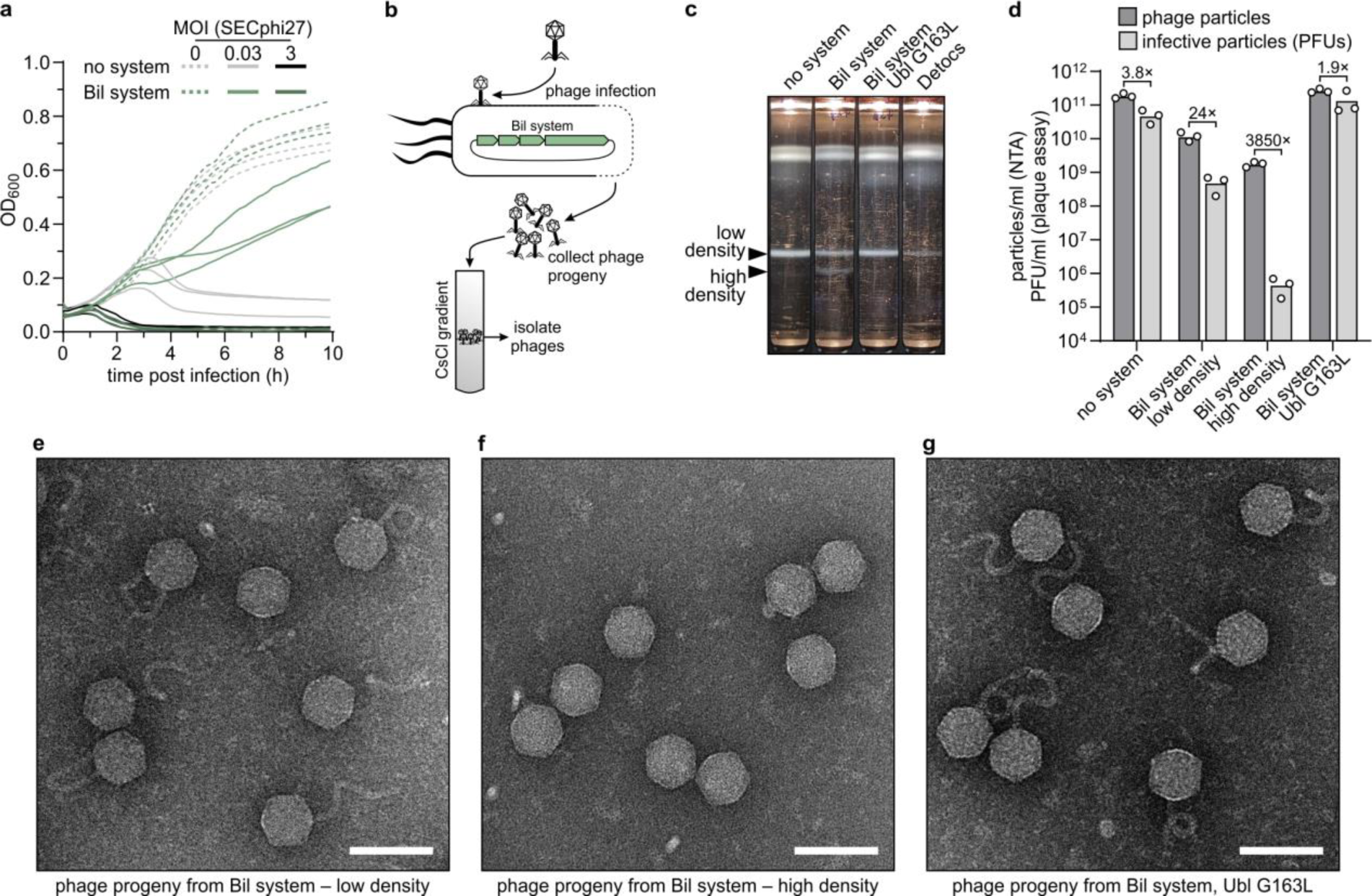
The Bil system causes the production of non-infective phage particles. **a,** Growth curves of *E. coli* cells expressing a control plasmid or the Bil system, either uninfected or infected with phage SECphi27 at a multiplicity of infection (MOI) of 0.03 or 3. Three independent replicates are shown as individual curves. **b,** Overview of the phage purification process. Bacteria expressing the Bil system were infected with phage SECphi27 at an MOI of 0.1 for 3 h to allow at least two full rounds of phage replication. The resulting lysate was filtered to obtain the phages, which were further purified using CsCl gradient ultracentrifugation. **c,** Photographs of CsCl gradients loaded with phages obtained from bacteria expressing the indicated systems. Only the wild-type Bil system leads to the appearance of a second, higher density band of phages. **d,** Particle concentration and number of plaque-forming units (PFUs) of phages obtained as shown in panel **c**, measured by nanoparticle tracking analysis and plaque assays, respectively. Bars represent the average of three independent replicates with individual data points overlaid. Fold difference between number of particles (measured via NTA) and number of infective particles (PFUs) is presented for each condition. **e-g,** Representative negative staining transmission electron microscopy images of phages obtained from the low-density (**e**) or high-density (**f**) bands of bacteria expressing the wild-type Bil system or the band from bacteria expressing the mutant Bil system (**g**). Scale bars represent 100 nm.

To examine whether phage progeny emerge from infected Bil-expressing cultures, we collected filtered supernatants from Bil-expressing cells that were lysed following infection with SECphi27. We then used isopycnic CsCl gradient centrifugation to achieve a concentrated phage preparation (Figure 2B). Mature phages concentrated in a CsCl gradient typically localize to a single band where the density of phage particles matches the local density of the gradient^21^. However, supernatant derived from infected Bil-expressing cells formed two distinct bands on the CsCl gradient (Figure 2C). One of these bands was at the same density in the gradient as the band derived from phages propagated on control cells, suggesting that this band represents mature phages. The second band ran lower in the gradient and hence represented particles of higher density. The second band was not observed for phages propagated on a mutant Bil system, suggesting that generation of higher-density particles is a property of a functional Bil system (Figure 2C).

We next quantified the phages isolated from each band in the CsCl gradient using nanoparticle tracking analysis (NTA), a biophysical technique that can count particles with diameters of 10-1000 nm by tracking their Brownian motion^22^. Using the same phage samples, we quantified the number of infective particles (plaque forming units, PFUs) using standard plaque assays on an indicator strain. For phages propagated on control cells or on cells expressing the mutant Bil system, the number of particles measured by the biophysical NTA technique was largely in agreement with the measured PFUs, confirming that the NTA method is suitable for accurately counting virions^23^ (Figure 2D). By contrast, the number of particles measured in the unique high-density band derived from cells expressing the Bil system was ∼4000-fold greater than the PFUs measured for the same sample, indicating that over 99.9% of particles in this band were non-infective (Figure 2D). Similar measurements for the low-density band derived from infected Bil-expressing cells, which presumably includes mature phages, showed that this band contained ∼24 times more particles measured by NTA than PFUs, suggesting that even in the low-density band, the majority of particles are non-infective (Figure 2D).

To further characterize the phages emerging from bacteria expressing the Bil system, we performed negative staining transmission electron microscopy on particles derived from each of the two bands of the CsCl gradient. Phages from the low-density band were morphologically indistinct from normal phages, possessing tails and DNA-filled capsids typical for the SECphi27 siphovirus (Figure 2E). By contrast, particles from the high-density band mostly consisted of head-only, DNA-filled capsids without a tail (Figure 2F), explaining why these particles were not infective (Figure 2D). Consistent with this observation, our NTA measurements showed that particles from the high-density band were of substantially smaller sizes (Figure S2). This observation explains why these tailless phages form a denser band in the CsCl gradient: the removal of the tail increases the ratio of DNA to protein within the particle, which in turn increases its buoyant density^24,25^. As expected, phages propagated on the mutated version of the Bil system showed normal morphology (Figure 2G). These results show that the Bil system somehow interferes with phage assembly, causing the generation of tailless phages as well as tailed phages with reduced infectivity.

To rule out the possibility that the production of partially assembled phages is a general property of anti-phage defense systems that protect against SECphi27, we propagated SECphi27 on *E. coli* cells expressing Detocs, a defense system that depletes cellular ATP in response to infection, thus aborting the phage replication cycle^26^. Propagation of SECphi27 on Detocs-expressing cells yielded less phage progeny, consistent with previous observations that Detocs protects against this phage^26^ (Figure 2C). However, SECphi27 progeny obtained from Detocs-expressing bacteria ran as a single band on the CsCl gradient, and the denser band observed for phages derived from the Bil system was not visible for Detocs-derived phages (Figure 2C). These results further support that the production of tailless phages is a unique property of the Bil defense system.

### The central tail fiber of the phage is the target of the Bil system

While it is obvious why tailless phages would be non-infective, it remained unclear why Bil-derived phages from the low-density CsCl band, which seemed properly assembled and tailed according to electron microscopy examination, still exhibited reduced infectivity (Figure 2D, E). Based on the properties of the Bil system, we hypothesized that these phages might have been conjugated to the Bil Ubl protein in a way that interferes with their infectivity. To test this hypothesis, we experimented with a Bil system variant in which the BilA Ubl protein was N-terminally fused to an HA-tag. This tagged system showed defense against phage SECphi27, indicating that N-terminal tagging of the Ubl protein does not interfere with the function of the Bil system (Figure S3A). We then propagated phages on cells expressing the tagged Bil system, isolated these phages using CsCl gradients, and immunoprecipitated the phages using anti-HA antibodies. Western blot analysis of proteins from pulled-down phages revealed a major band at ∼150 kDa, indicating a single phage protein conjugated to the Ubl (Figure 3A). Only one protein of SECphi27, the central tail fiber, is large enough to generate a protein band of this size (central tail fiber: ∼132 kDa; HA-Ubl: ∼19 kDa). Mass spectrometry analysis identified the SECphi27 central tail fiber as well as the Ubl protein in this band (Table S1), showing that the ∼150 kDa band indeed represents a Ubl∼central tail fiber adduct. These results suggest that the Bil system conjugates its Ubl to the phage central tail fiber.

**Figure 3.**
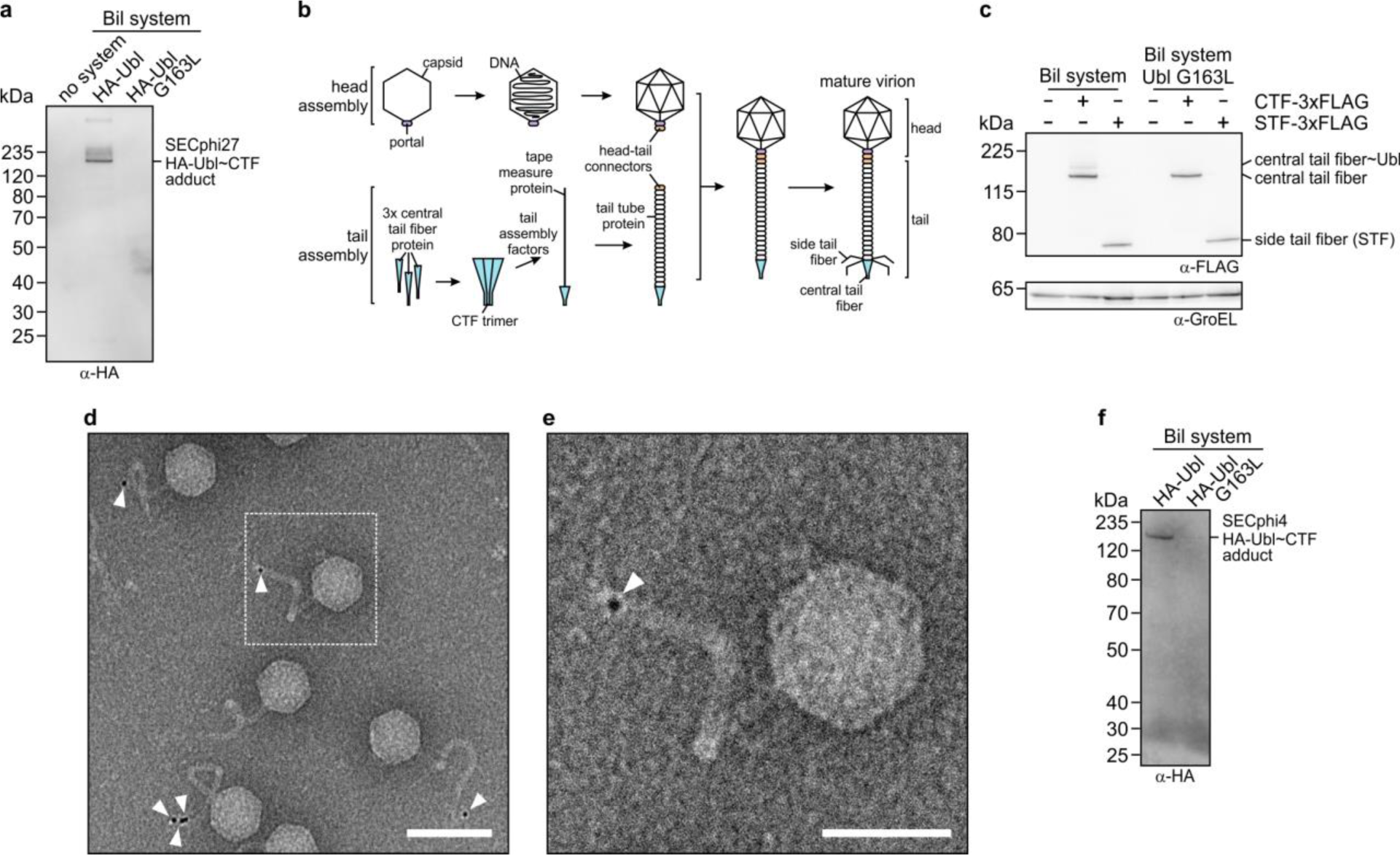
The phage central tail fiber is the target of the Bil system. **a,** Immunoprecipitation of HA-tagged Ubl protein from SECphi27 phages collected following infection of bacteria expressing wild-type and mutant Bil system, analyzed by western blotting. Only one phage protein, the CTF of SECphi27, is revealed to be conjugated to HA-Ubl. **b,** Simplified representation of the siphophage virion assembly pathway, based on studies of phage λ^13,27,28^. **c,** Co-expression of the Bil system with either the central tail fiber (CTF) or the side tail fiber (STF) of SECphi27, analyzed by western blotting. GroEL was used as loading control. **d,** Representative immunogold labeling transmission electron microscopy image of SECphi27 phages isolated from bacteria expressing the HA-Ubl Bil system. White arrowheads represent gold labeling of HA-Ubl. Scale bar represents 100 nm. See Figure S4 for additional images. **e,** Magnification of the dashed area in panel **d**. White arrowhead represents gold labeling of HA-Ubl. Scale bar represents 50 nm. **f,** Immunoprecipitation of HA-tagged Ubl protein from SECphi4 phages collected following infection of bacteria expressing wild-type and mutant Bil system, analyzed by western blotting. Only one phage protein, the CTF of SECphi4, is revealed to be conjugated to HA-Ubl.

The central tail fiber (CTF, also called spike protein or tip attachment protein) is an essential structural component of many siphophages, forming the tip of the tail in the mature virion (Figure 3B). The CTF contains the receptor-binding domain of the phage, which is responsible for host receptor recognition^14^. Furthermore, the CTF is essential for assembly of the phage tail^13,27,28^. In phage λ, for which the tail assembly cascade was studied in detail, it was shown that tail assembly begins from trimerization of the CTF. The CTF trimer then recruits tail assembly factors, which in turn recruit the tail tape measure protein, initiating the polymerization of the tail tube proteins. Once the tail is mature, it is attached to the DNA-filled capsid, which is assembled in the cell independently of the tail, to ultimately produce the mature tailed virion^13,27,28^ (Figure 3B). Therefore, conjugation of a Ubl protein to the CTF might interfere with assembly of the phage tail, explaining why many of the virions emerging from infected Bil-expressing cells are tailless. In support of this hypothesis, pulldown attempts of the fraction of tailless phages (high-density CsCl band) that were propagated on the HA-Ubl Bil system did not retrieve any tagged proteins (Figure S3B), since the CTF is missing from these tailless phages.

To confirm that the Bil system specifically targets the phage CTF, we co-expressed a 3xFLAG-tagged version of the CTF together with the Bil system in the absence of other phage components, and then analyzed whole cell lysates via western blotting. This revealed two bands: one corresponding to the unmodified CTF, and one to the Ubl-CTF adduct (Figure 3C). In contrast, when we expressed the CTF with the mutant Bil system, only the unmodified CTF could be observed (Figure 3C). To further examine whether the Bil system is specific to the CTF, we performed the same experiment with the side tail fiber of the phage, which is also localized at the distal part of the tail (Figure 3B), but could not observe conjugation of the Ubl to the side tail fiber (Figure 3C). These results further substantiate that the Bil system specifically conjugates the Ubl to the phage CTF protein.

To further test whether the fraction of fully assembled, tailed phages that emerged from infected Bil-expressing cells are specifically modified by the Ubl on their CTF, we performed immunogold labeling transmission electron microscopy experiments. In these experiments, phages propagated on a Bil system with an HA-tagged Ubl were probed with anti-HA antibodies and then with secondary antibodies attached to 6 nm colloidal gold particles. In agreement with our biochemical data, gold labeling was observed exclusively at the tip of the phage tails — the position of the CTF (Figures 3D, E, S4). We occasionally noted two gold particles, and sometimes more, at the same tail tip, possibly representing Ubl conjugation on two or three monomers of the CTF trimer, or representing multiple secondary antibodies bound to the same anti-HA primary antibody (Figure S4). The same experiment with a tagged but mutated Ubl protein did not lead to labeling of the tail tip (Figure S5A). These results demonstrate that the Bil system attaches the Ubl protein to the CTF protein of the phage.

The Bil system defends against phages spanning multiple different taxonomical families^12^. To test whether the system protects against diverse phages via the same mechanism, we selected SECphi4, a siphophage from the *Dhillonvirus* genus whose genome (∼45 kb) has no detectable sequence similarity to the genome of SECphi27 (∼52 kb) and against which the Bil system strongly protects (Figure S5B). We purified SECphi4 phages propagated on a strain expressing the HA-Ubl-tagged Bil system and immunoprecipitated these phages with anti-HA antibodies. Western blot analysis of proteins from immunoprecipitated phages revealed only one SECphi4 protein that was conjugated to the Ubl (Figure 3F). Mass spectrometry analysis identified this protein band as an adduct of the CTF of SECphi4 and the Ubl of the Bil system (Table S2). In support of this observation, co-expression of a tagged version of the CTF of SECphi4 with the Bil system in the absence of phage infection showed that the system is able to conjugate the Ubl protein to the SECphi4 CTF (Figure S5C). These results demonstrate that the Bil system functions similarly when protecting against distantly related phages, and that the phage CTF is the target of the Bil system for Ubl conjugation.

While SECphi27 and SECphi4 have no sequence similarity on the nucleotide level, 16 of the 85 (∼19%) predicted proteins in the SECphi27 proteome show homology to proteins of SECphi4 (Table S3). Notably, 11 of these 16 proteins are predicted tail structure proteins or tail assembly proteins, including the CTF, suggesting that these two phages possess similar tail structures (Table S3). The CTF proteins of SECphi4 and SECphi27 show ∼34% sequence identity on the amino acid level, and prediction of their protein structures using AlphaFold2 revealed substantial structural similarities between the two proteins (Figure S5D), which could explain how the Bil system may recognize both proteins and defend against phages from different families. The fact that two distantly related phages that share little but their tail structures are both targeted by the Bil system further supports that the primary mechanism of the Bil system involves interfering with phage tail assembly by conjugating a Ubl protein on the phage CTF.

## Discussion

In this study, we investigated the mechanism of immunity provided by a bacterial defense system capable of conjugating an ISG15-like Ubl protein. Our data support a model in which this system specifically conjugates the ubiquitin-like protein to the central tail fiber of the phage (Figure 4). This conjugation can prevent tail assembly and lead to the generation of non-infective tailless phages or produce infection-impaired tailed phages whose tail tip is obstructed by a covalently attached Ubl. This mode of action does not save the infected cell from eventually being lysed by the phage, but since the particles that emerge from the lysed cell are mostly non-infective, the Bil system ends up protecting the surrounding bacteria from the spread of phage epidemic (Figure 4). The mechanism of the Bil system is different than that of abortive infection (Abi) systems, which actively kill the infected cells once phage infection is detected^29,30^. However, a similar outcome of culture-level protection is achieved by both Bil and Abi systems.

**Figure 4.**
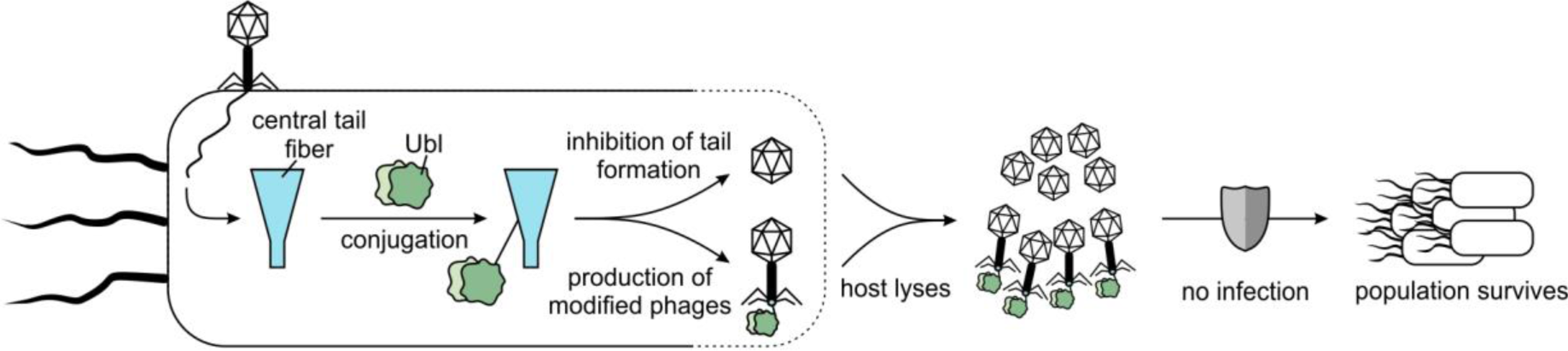
Model for the mechanism of action of the Bil defense system. After phage infection, the Bil system conjugates its Ubl protein to the central tail fiber protein of the phage, which leads either to the inhibition of tail formation and generation of tailless phages or to the production of modified phages with obstructed tail tips. After the host bacterium is lysed by the phage, these two populations of phages are released but are unable to infect the surrounding bacteria.

While our results reveal the CTF as the target of the Bil system, the mechanism of CTF recognition remains to be studied. Our observations using two taxonomically distant phages suggest that a structural motif in the CTF may be the determining factor of recognition (Figure S5D). The Bil system also defends against T5^12^, a siphophage with a tail structure different than that of the two phages studied here^31^. Nevertheless, there are substantial structural similarities between the N-terminal half of the SECphi27 CTF and the T5 baseplate hub protein pb3, a central structural protein of the T5 tail tip (Figure S6A). The baseplate hub protein of T5 was shown to have structural similarity with the baseplate hub of phage T4^31^ (Figure S6B), which might explain how the Bil system protects against T-even phages as well^12^.

An aspect of the Bil system that was not studied here is the role of the deubiquitinase (DUB) protein. This protein is essential for the proper activity of the system, as disruption of its active site renders the system inactive (Figure 1D). It is possible that the DUB removes the Ubl from proteins when nonspecific conjugations occur in order to keep a pool of free Ubl until phage infection. This could also prevent potential toxic effects of nonspecific Ubl conjugation to bacterial proteins. In possible support for this hypothesis, we found that the DUB of the *Collimonas* Bil system studied here is able to cleave the Ubl when it is fused to GFP (Figure S7). Interestingly, some siphophages encode tail assembly proteins containing DUB domains whose functions in phage tail assembly are not currently understood^32^. Since the Bil system interferes with phage tail assembly, it is tempting to speculate that the inclusion of DUB domains in phage tail assembly proteins is an anti-defense measure to protect the phage from the action of the Bil system by cleaving the Ubl off the central tail fiber.

Proteins with E1, E2 and DUB domains are also present as accessory proteins in the anti-phage defense systems CBASS and Pycsar^33–35^. These systems depend on second messenger signaling molecules that are produced in response to phage detection, and it was shown that the proteins that generate the second messenger molecules can be conjugated by the E1 and E2 enzymes to unknown targets via a ubiquitin-like mechanism, thereby priming their enzymatic activity^36,37^. In contrast to the Bil system, E1/E2-dependent conjugation in CBASS is only essential for defense against some phages, while dispensable for others^33,36,37^. Since the Ubl of the Bil system has no predicted enzymatic activity^12^, it is likely that the primary function of the Bil system is to interfere with phage assembly and cause the generation of non-infective particles. It is possible that conjugation of CBASS proteins to phage structural components may be another aspect of CBASS defense.

The Bil system we studied is a representative of a large family of defense systems encoding E1, E2 and Ubl proteins^12^. In some of these systems, the Ubl contains a single ubiquitin-like domain (instead of the two fused domains in the Bil system), and in some cases, the gene with the DUB domain is missing^12^. It remains to be determined whether all these systems target the phage CTF, or whether some of them may target other components of the phage.

The Bil system can be viewed as analogous to the human ISG15 system, in the sense that both systems encode a double ubiquitin Ubl and dedicated Ubl conjugation machinery, and that both systems protect against viruses^6,9,10^. Despite extensive studies over the course of several decades, the mechanism of ISG15 antiviral defense is not entirely understood^6,9,10^. A variety of cellular and viral targets were suggested for ISG15^6^, and it was also suggested that the dedicated E1, E2 and E3 enzymes of ISG15 non-specifically conjugate it to newly synthesized proteins in infected cells to interfere with viral reproduction^38^. Our discovery that the bacterial analog of the human ISG15 interferes with phage assembly by targeting viral structural proteins may guide future studies on the human counterpart.

## Methods

### Bacterial strains and phages

The Bil system (IMG^39^ gene IDs: BilA: 2609810443; BilB: 2609810442; BilC: 2609810441; BilD: 2609810440) and its mutant variants were expressed in *E. coli* MG1655. A plasmid containing *gfp* instead of the Bil system was used as negative control throughout this study (denoted “no system”). Bacteria were grown in MMB (LB supplemented with 0.1 mM MnCl_2_ and 5 mM MgCl_2_) at 25°C with shaking at 200 rpm, and the appropriate antibiotics (100 µg/ml ampicillin and/or 30 µg/ml chloramphenicol) were added. The bacterial strains and phages used in this study are listed in Table S4. Infection was performed in MMB with or without 0.5% agar.

### Plasmid and strain construction

All plasmids with their respective inserts used in this study are listed in Table S5. Variants of the Bil system were generated by amplification of the plasmid containing the wild-type Bil system^12^ using primers that contained the desired modification followed by treatment with KLD enzyme mix (NEB) according to the manufacturer’s instructions to obtain transformable plasmids. Plasmids encoding phage proteins were constructed by amplification of the desired genes from the phage genomes and assembly into the pBbA6c backbone using NEBuilder HiFi DNA Assembly Master Mix (NEB) according to the manufacturer’s instructions. After sequence verification, plasmids were transformed into *E. coli* MG1655 using TSS transformation^40^.

### Plaque assays

To test the defensive properties of the Bil system and its mutant versions, the small drop plaque assay was used^41^. 300 μl of an overnight culture of bacteria containing the respective version of the Bil system or the negative control strain were mixed with 30 ml of melted 0.5% MMB agar containing 0.2% arabinose. The mixture was poured into a 10 cm square plate and left to solidify for 1 hour at room temperature. Ten-fold serial dilutions of phages in MMB were prepared and 10 μl drops of each dilution were plated on the bacterial layer. The plates were incubated overnight at 25°C. Plaque-forming units (PFUs) were counted after overnight incubation.

### Liquid infection assays

Overnight cultures of bacteria were diluted 1:100 in MMB and incubated until they reached an OD_600_ of 0.3, after which 180 μl of the culture was transferred into a 96-well plate containing 20 μl of either MMB (for the uninfected control) or phage lysate for a final MOI of 3 or 0.03. Plates were incubated at 25°C with shaking in a Tecan Infinite 200 plate reader and the OD_600_ was measured every 10 min. Infections were performed in biological triplicates in three separate plates with technical triplicates in each plate. Three wells were filled with medium in each plate to serve as the blank, which was subtracted from the OD_600_ values of the wells containing bacteria.

### Phage purification

To purify phages for electron microscopy and nanoparticle tracking analysis, 25 ml cultures of bacteria were grown with the appropriate antibiotics and 0.2% arabinose at 25°C with shaking until they reached an OD_600_ of 0.55. Then, the cultures were infected with phages at an MOI of 0.1, followed by incubation at 25°C for 3 h with shaking to allow for at least two full rounds of phage replication. To clear the cultures of unlysed bacteria and debris, they were centrifuged at 3,300 *g* and 4°C for 10 min and filtered through a 0.2 µm filter. 12.5 ml of 20 mM Tris-HCl, pH 7.5, 3 M NaCl, 30% PEG-8000 was added to the cleared phage lysates to precipitate the phages overnight at 4°C. The precipitated phages were collected by centrifugation at 25,000 *g* and 4°C for 30 min, followed by decanting of the supernatant and centrifugation at 10,000 *g* and 4°C for 2 min. The remaining supernatant was removed and the phage pellets were soaked in 1 ml of 50 mM Tris-HCl, pH 7.5, 150 mM NaCl, 10 mM MgSO_4_ on ice for 30 min to loosen them. The phage pellets were finally dissolved in the added buffer by careful pipetting and loaded on CsCl step gradients (1 ml of ρ = 1.3, 4 ml of ρ = 1.4, 4 ml of ρ = 1.5, 1 ml of ρ = 1.7; all in 50 mM Tris-HCl, pH 7.5, 10 mM MgSO_4_) formed in open-top polyclear ultracentrifugation tubes (Seton Scientific). The gradients were centrifuged in an SW41 rotor (Beckman) at 25,000 rpm and 4°C for 3 h and the phage bands collected by needle side-puncture. The extracted phages were dialyzed against 20 mM Tris-HCl, pH 7.5, 150 mM NaCl, 10 mM MgSO_4_ overnight at 4°C using Pur-A-Lyzer Maxi 12000 dialysis tubes (Sigma-Aldrich). The phages were further concentrated by centrifugation in a TLA-110 rotor (Beckman) at 45,000 rpm and 4°C for 1 h. After removal of the supernatant, the phages were soaked in 50 µl of 20 mM Tris-HCl, pH 7.5, 150 mM NaCl, 10 mM MgSO_4_ on ice for 30 min, followed by careful resuspension to obtain the final samples.

### Immunoprecipitation

For immunoprecipitation of HA-tagged Ubl from whole cell lysates (Figure 1E), 50 ml of each strain was grown at 25°C to an OD_600_ of 0.3 with 0.2% arabinose, followed by centrifugation at 3,300 *g* and 4°C for 10 min to pellet the bacteria. The supernatant was discarded and the pellet frozen at –80°C. The pellet was thawed on ice and dissolved in 750 µl of TBS-T (20 mM Tris-HCl, pH 7.5, 150 mM NaCl, 0.05% Tween 20), followed by transfer to a 2 ml tube with lysing matrix E (MP Biomedicals) and mechanical lysis using a FastPrep-24 instrument (MP Biomedicals) at 6 m/s and 4°C for 40 s. The lysate was cleared via centrifugation at 13,000 *g* and 4°C for 10 min. 25 µl of anti-HA magnetic beads (Thermo Scientific) was washed with 1 ml of ice-cold TBS-T and the cleared lysate was added to the washed beads, followed by rotation at 4°C for 1 h to allow binding of HA-Ubl to the beads. The lysate was removed and the beads washed three times with 500 µl ice-cold TBS-T for 1 min. The beads were washed with 500 µl of water and finally resuspended in 35 μl 1x Bolt LDS Sample Buffer (Thermo Scientific), 20 µl of which was subjected to non-reducing SDS-PAGE.

For immunoprecipitation of HA-tagged Ubl from isolated phages (Figures 3A, F and S3B), phages were purified as described above, except that the final concentration step after dialysis was skipped. Phages were added to 25 µl of washed beads and immunoprecipitation performed as described above. The beads were resuspended in 35 μl 1x Bolt LDS Sample Buffer supplemented with 50 mM DTT, of which 10 µl was used for western blotting and 20 µl was subjected to SDS-PAGE and mass spectrometry.

### Analysis of covalent, thioester-based complex formation between the Ubl and E1/E2 proteins

To analyze covalent thioester-based complexes between the Ubl and E1/E2 proteins (Figure S1A, C), 10 ml of each strain was grown to an OD_600_ of 0.5 or 0.8 with 0.2% arabinose to express the Bil system. An equivalent to 0.1 OD_600_ of cells was collected and centrifuged at 13,000 *g* and 4°C for 2.5 min to pellet the bacteria. The pellet was resuspended in 100 µl 1x Bolt LDS Sample Buffer, which was split into two tubes. One of the tubes was supplemented with 50 mM DTT to achieve reducing conditions able to break potential thioester bonds^18^. 10 µl each of the reduced and the non-reduced samples were subjected to western blotting.

### Co-expression analysis

For co-expression analyses (Figures 3C, S5C and S6C), 10 ml of each strain was grown to an OD_600_ of 0.5 with 0.2% arabinose to express the Bil system or the Ubl-GFP fusion protein. Then, the phage CTFs/STFs or the DUB were induced with 0.1 mM IPTG or 50 ng/ml aTc, respectively, for 30 min. An equivalent to 0.1 OD_600_ of cells was collected and centrifuged at 13,000 *g* and 4°C for 2.5 min to pellet the bacteria. The pellet was resuspended in 100 µl 1x Bolt LDS Sample Buffer supplemented with 50 mM DTT, of which 10 µl were subjected to western blotting.

### Protein gel electrophoresis and western blotting

For protein analysis, the indicated amounts of protein sample were boiled at 95°C for 3 min and separated by 4–12% Bis-Tris SDS-PAGE (Thermo Scientific) either in 1x MES buffer (Thermo Scientific) or 1x MOPS buffer (Thermo Scientific) to resolve smaller and larger proteins, respectively, at 200 V. Coomassie staining was performed by addition of InstantBlue Coomassie Protein Stain (Abcam) for 1 h, followed by destaining with water. For western blotting, the gel was transferred to a PVDF membrane (Thermo Scientific) for 1 hour at 20 V in 1x transfer buffer (Thermo Scientific) and probed with the appropriate primary antibody diluted in TBS-T with 3% BSA: rabbit anti-HA (Sigma Aldrich #H6908; 1:3,000 dilution), mouse anti-FLAG (Sigma Aldrich #F1804; 1:10,000 dilution), mouse anti-GFP (Sigma Aldrich #11814460001; 1:5,000 dilution) or HRP-conjugated Strep-Tactin (IBA #2-1502-001; 1:50,000 dilution). Visualization of the primary antibody was performed using HRP-conjugated goat anti-rabbit secondary antibody (Thermo Scientific #31460; 1:10,000 dilution) or HRP-conjugated goat anti-mouse secondary antibody (Thermo scientific #31430; 1:10,000 dilution) and incubation with ECL solution (Merck Millipore). No secondary antibody was used in the case of the HRP-coupled Strep-Tactin. Where appropriate, the membranes were stripped using Restore PLUS Western Blot Stripping Buffer (Thermo Scientific) and probed with primary rabbit anti-GroEL antibody (Sigma Aldrich #G6532) as loading control.

### Mass spectrometry

To identify the phage proteins observed to be conjugated to HA-Ubl, SDS-PAGE was run and stained with Coomassie as described above. The band of interest was cut from the gel and the same area of the gel was cut for the control as well. The gel bands were subjected to in-gel tryptic digestion. The resulting peptides were analyzed using nanoflow liquid chromatography (nanoAcquity) coupled to high resolution, high mass accuracy mass spectrometry (Q Exactive HF). Each sample was analyzed on the instrument separately in a random order in discovery mode. The data was searched against the *E. coli* K-12 database appended with the SECphi27, SECphi4 and Bil system proteomes, as well as common lab contaminants. Carbamidomethylation of C was set as a fixed modification, while oxidation of M and deamidation of NQ were defined as variable ones.

### Negative staining transmission electron microscopy

For negative staining, formvar/carbon-coated nickel grids (300 mesh, Electron Microscopy Sciences) were glow discharged in an Evactron CombiClean Decontaminator (XEI Scientific) for 1 min at 0.53 mbar (air) and 18 W. 3 µl of phage sample was directly applied to the grid, incubated for 1 min and then blotted with filter paper (Whatman Grade 1). The grid was briefly washed on a drop of water and blotted with filter paper. Staining was performed by touching a 15 µl drop of staining solution (freshly made 2% ammonium molybdate, pH 6.5), followed by blotting with filter paper. This step was repeated once and then the grid was floated on top of a 15 µl drop of staining solution for 1 min. Finally, the grid was blotted with filter paper and air dried. Imaging was performed using a Tecnai T12 transmission electron microscope (Thermo Fisher Scientific) at accelerating voltage of 120 kV, with a TVIPS TemCam-XF416 CMOS camera and SerialEM acquisition software^42^.

### Immunogold labeling transmission electron microscopy

For immunogold labeling, a previously published protocol was followed with some changes^43^. Formvar/carbon-coated nickel grids (300 mesh) were glow discharged as above. 3 µl of phage sample was directly applied to the grid, incubated for 1 min and then blotted with filter paper. The grid was briefly washed on a drop of water, blotted with filter paper and floated on a 20 µl drop of blocking buffer (PBS with 0.1% Tween 20 and 0.3% IgG-free BSA) for 30 min in a closed, humidified chamber. The grid was blotted with filter paper and floated on a 20 µl drop of primary anti-HA antibody (Sigma-Aldrich #H6908) diluted 1:500 in blocking buffer for 1 h in a closed, humidified chamber. Following blotting with filter paper, the grid was washed five times by floating it on a 20 µl drop of wash buffer (PBS with 0.1% Tween 20 and 0.03% IgG-free BSA) for 3 min and blotted with filter paper after each wash. Next, the grid was floated on a 20 µl drop of anti-rabbit secondary antibody coupled to 6 nm colloidal gold particles (Electron Microscopy Sciences #25104) diluted 1:50 in blocking buffer for 1 h in a closed, humidified chamber. The grid was washed as before, briefly washed twice on drops of water, blotted with filter paper and stained as described above. Imaging was performed as described above.

### Nanoparticle tracking analysis

For nanoparticle tracking analysis (NTA), phages were purified as described above, except that the final concentration step after dialysis was skipped and the volumes of the phage samples was increased to 1.25 ml by addition of buffer (20 mM Tris-HCl, pH 7.5, 150 mM NaCl, 10 mM MgSO_4_). 1 ml of each phage sample (negative control: diluted 1:100; Bil system low-density band: diluted 1:10; Bil system high-density band: undiluted; mutated Bil system: diluted 1:100) was injected into the flow cell of a NanoSight NS300 instrument (Malvern Panalytical) equipped with a 405 nm laser. Measurements were performed using the NanoSight NTA 3.4 software with the following parameters: the camera level was set to 10 and for each sample a standard measurement with 3 captures of 60 s per sample was run. The captured data was analyzed and exported using the default parameters of the standard measurement protocol with a detection threshold of 2. To obtain the final numbers of phage particles per ml, the reported values were multiplied with the respective dilution factors of each sample. The same phage samples were used to measure PFUs using plaque assays as described above, except that a wild-type *E. coli* strain without plasmids was used as the indicator strain and no arabinose was added to the medium.

### Structural predictions

Structural predictions of proteins and complexes was performed using AlphaFold2^15^ and AlphaFold-Multimer^19^, respectively. The predictions were run via ColabFold v1.5.2^44^ using default parameters. To compare the structures of proteins/domains and calculate the respective root mean square deviation (RMSD) values, the best-ranking structure predictions were chosen and aligned using the “super” method in the alignment plugin of PyMOL 2.5.5^45^. To visualize these aligned structures, they were exported from PyMOL in their aligned states and imported into UCSF ChimeraX 1.6.1^46^, where the structure images were generated.

### Sequence conservation analysis

To analyze conserved residues, non-redundant BilA homologs collected in a previous study^12^ were aligned using Clustal Omega^47^ using default parameters. Sequence logos were generated using WebLogo (https://weblogo.berkeley.edu/logo.cgi). 12 of the aligned proteins^12^ (Table S6) contained short (2–9 aa) overhangs downstream of the conserved C-terminal glycine, which are not shown in the sequence logo for clarity.

To compare the proteomes of SECphi27 (NCBI taxonomy ID: 2496550) and SECphi4 (NCBI taxonomy ID: 2729542), an all-versus-all blast analysis was performed using blastp^48,49^. Alignments with an E-value threshold of 1E-05 were considered as hits.

## Data availability

The mass spectrometry proteomics data have been deposited to the ProteomeXchange Consortium via the PRIDE partner repository^50^ with the dataset identifier PXD044622.

Plasmid inserts of the constructs used are available in Table S5.

## Acknowledgments

We thank the Sorek laboratory members for comments on earlier versions of this manuscript. We further thank Alon Savidor and Meital Kupervaser at the De Botton Protein Profiling Institute at the Weizmann Institute for help with mass spectrometry data generation and analysis. The electron microscopy studies were partially supported by the Irving and Cherna Moskowitz Center for Nano and BioNano Imaging (Weizmann Institute of Science). J.H. was funded by the Deutsche Forschungsgemeinschaft (DFG, German Research Foundation), grant 466645764 and a fellowship from the Council for Higher Education & Israel Academy of Science & Humanities (CHE/IASH) Excellence Fellowship Program for International Postdoctoral Researchers. R.S. was supported, in part, by the European Research Council (grant no. ERC-AdG GA 101018520), Israel Science Foundation (MAPATS Grant 2720/22), the Deutsche Forschungsgemeinschaft (SPP 2330, Grant 464312965), the Ernest and Bonnie Beutler Research Program of Excellence in Genomic Medicine, Dr. Barry Sherman Institute for Medicinal Chemistry, Miel de Botton, the Andre Deloro Prize, and the Knell Family Center for Microbiology.

## Author contributions

J.H. performed all experiments and visualized the data. S.G.W. performed electron microscopy. J.H. and R.S. conceptualized the study, analyzed the data and wrote the manuscript. All authors contributed to editing the manuscript and support the conclusions.

## Materials & correspondence

Correspondence and request for materials should be addressed to R.S.

## Supplementary Tables

**Table S1 – Mass spectrometry analysis of SECphi27 immunoprecipitation**

**Table S2 – Mass spectrometry analysis of SECphi4 immunoprecipitation**

**Table S3 – SECphi27 vs. SECphi4 proteome comparison**

**Table S4 – Bacterial strains and phages**

**Table S5 – Plasmids**

**Table S6 – BilA Ubl homologs**

**Figure S1.**
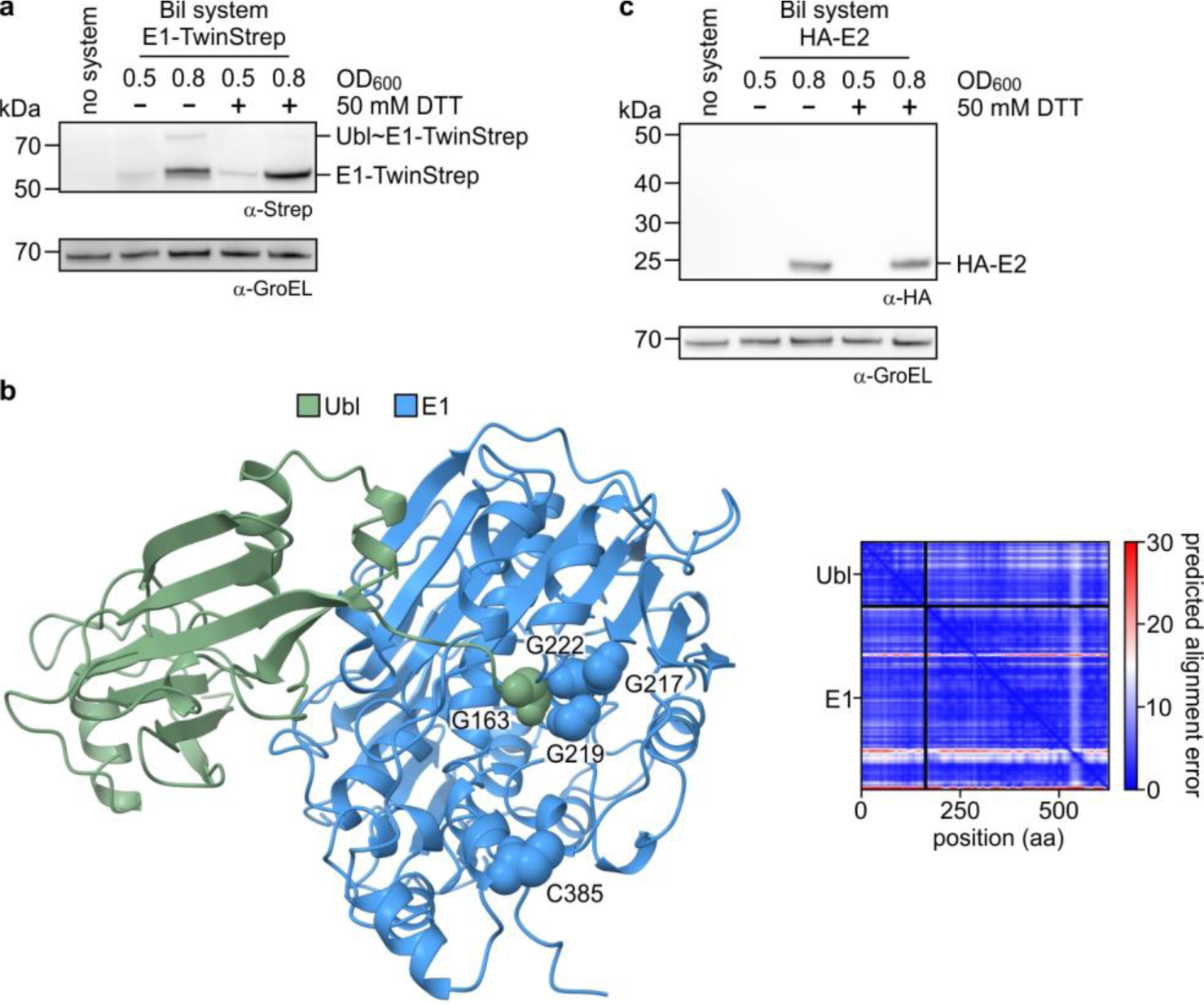
Interactions between the Bil Ubl and the E1 enzyme. **a,** Western blot of whole cell lysates of bacteria expressing the Bil system with a tagged E1. Two bands for the E1 are revealed: the ∼70 kDa band of which is reduced by DTT, indicating a thioester bond between the E1 and the Ubl. GroEL was used as loading control. Cells at OD600 of 0.8 are infected by phage SECphi27. **b,** AlphaFold-Multimer^19^ prediction of the interaction between the Ubl and the E1. A high confidence structure (model confidence of 0.94 and overall low predicted alignment error, right side of the panel) shows G163 (indicated as spheres) of the Ubl to be positioned at the nucleotide-binding loop of the E1 (G217, G219 and G222; indicated as spheres). **c,** Western blot of whole cell lysates of bacteria expressing the Bil system with a tagged E2. No covalent complex of the E2 could be observed. GroEL was used as loading control. Cells at OD600 of 0.8 are infected by phage SECphi27.

**Figure S2.**
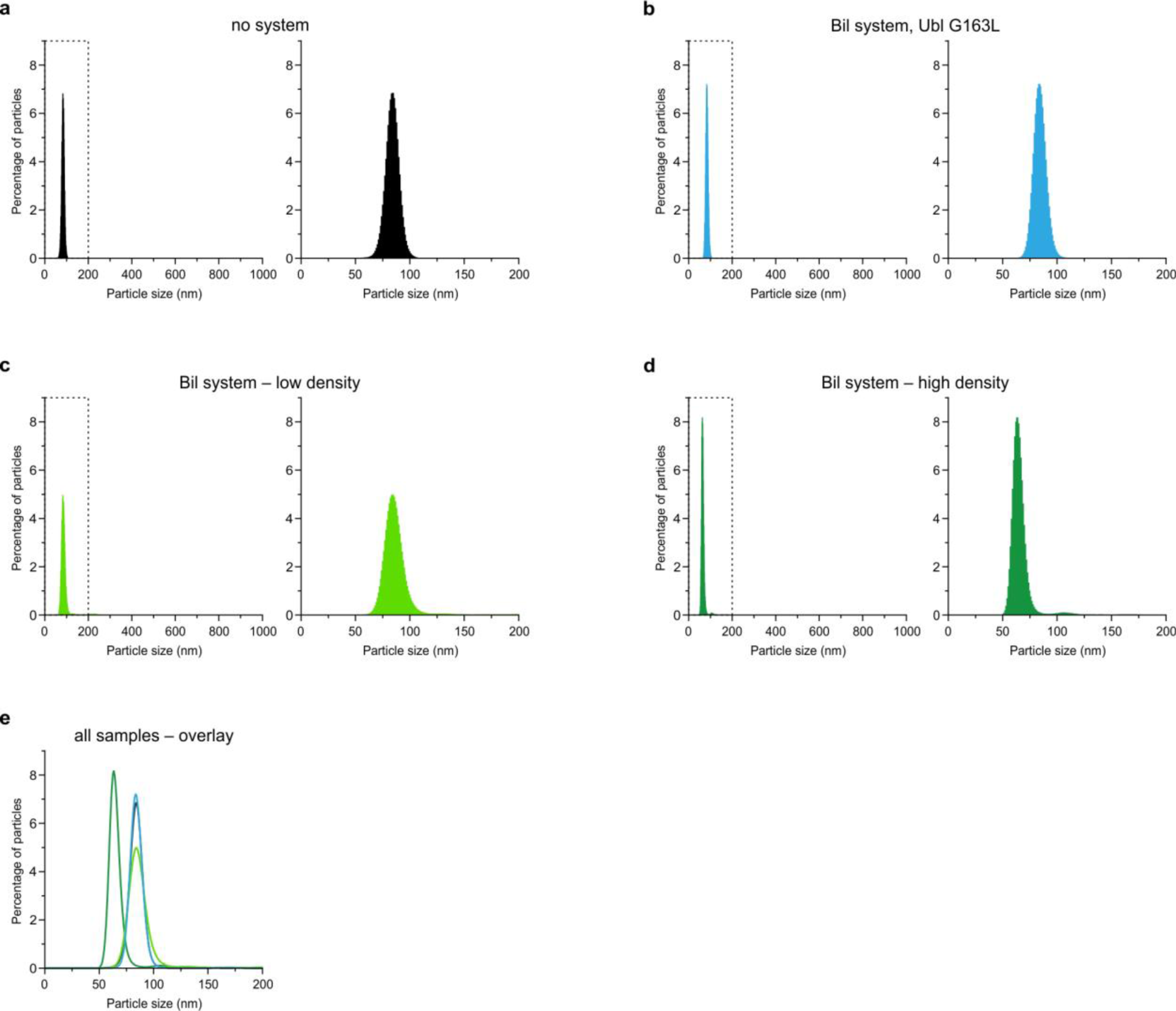
Phage particle size measurements. **a-d,** Particle size distributions of phages isolated from bacteria expressing the indicated systems, measured by nanoparticle tracking analysis. The dashed areas of each graph are magnified on the right of each graph. High and low density indicate phages isolated from high- and low-density bands on the CsCl gradient. **e,** Overlay of the particle size distributions of panels **a-d**.

**Figure S3.**
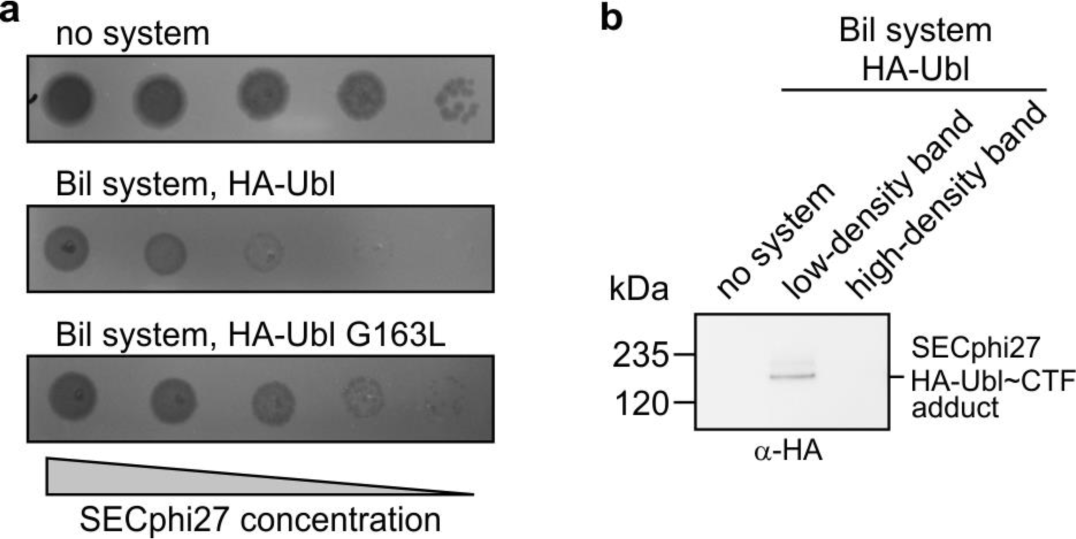
The phage central tail fiber is the target of the Bil system. **a,** Plaque assays showing that addition of an HA-tag to the Ubl of the Bil system does not interfere with phage defense. **b,** Immunoprecipitation of HA-tagged Ubl protein from SECphi27 isolated from bacteria expressing the Bil system, analyzed by western blotting. Phages from the low- and high-density bands of the CsCl gradient were immunoprecipitated separately.

**Figure S4.**
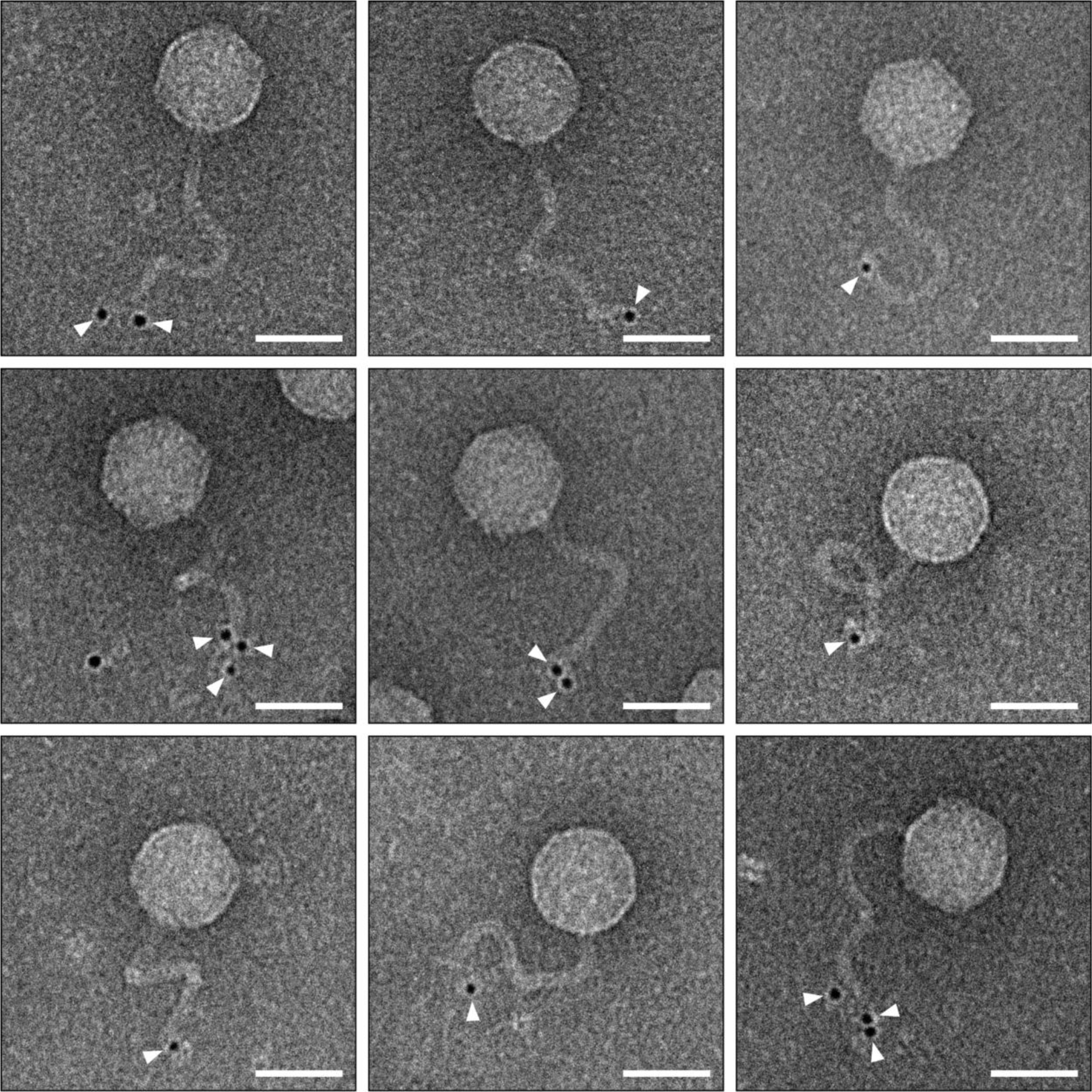
The Bil system modifies the tip of the phage tail. Immunogold labeling transmission electron microscopy images of SECphi27 phages isolated from bacteria expressing the HA-Ubl Bil system (related to Figure 3D). White arrowheads represent gold labeling of HA-Ubl. Scale bars represent 50 nm.

**Figure S5.**
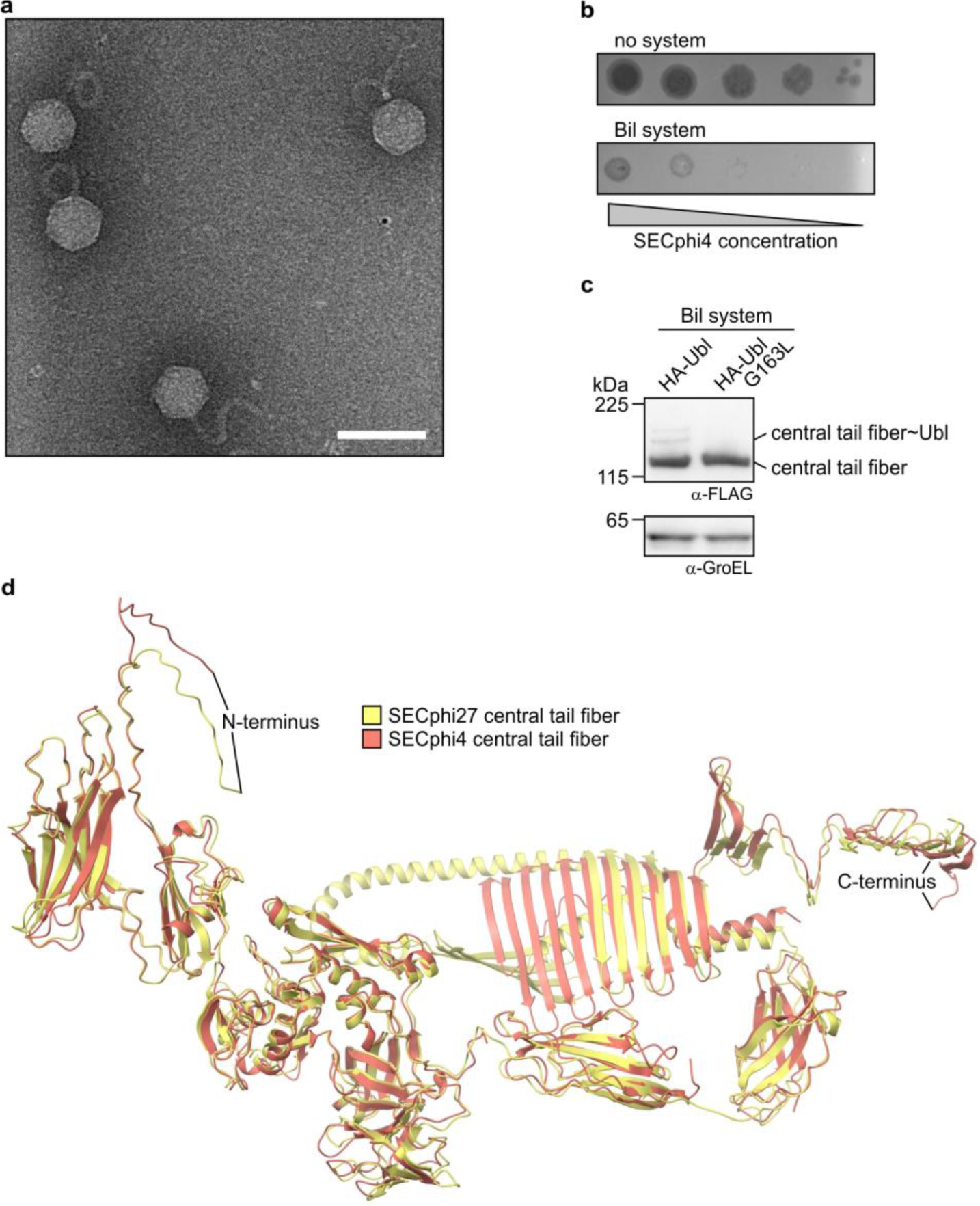
The central tail fiber is a conserved structure targeted by the Bil system. **a,** Representative immunogold labeling transmission electron microscopy image of SECphi27 phages isolated from bacteria expressing mutated HA-Ubl Bil system. No gold labeling could be observed on the phages. Scale bar represents 100 nm. **b,** Plaque assays showing the defense phenotype of the Bil systems against SECphi4. Data are representative of three independent experiments. **c,** Co-expression of the Bil system with the central tail fiber of SECphi4, analyzed by western blotting. GroEL was used as loading control. **d,** AlphaFold2^15^ predictions of the structures of the central tail fibers of SECphi27 (NCBI ID: YP_009965952.1) and SECphi4 (NCBI ID: QJI52569.1). The predicted structure of the SECphi4 central tail fiber was superimposed on the SECphi27 central tail fiber using the following domains: Residues 1–251 (RMSD = 0.91 Å), residues 252–628 (RMSD = 1.30 Å), residues 629–736 (RMSD = 1.11 Å), residues 737–829 (RMSD = 1.10 Å), residues 830–1139 (RMSD = 2.69 Å).

**Figure S6.**
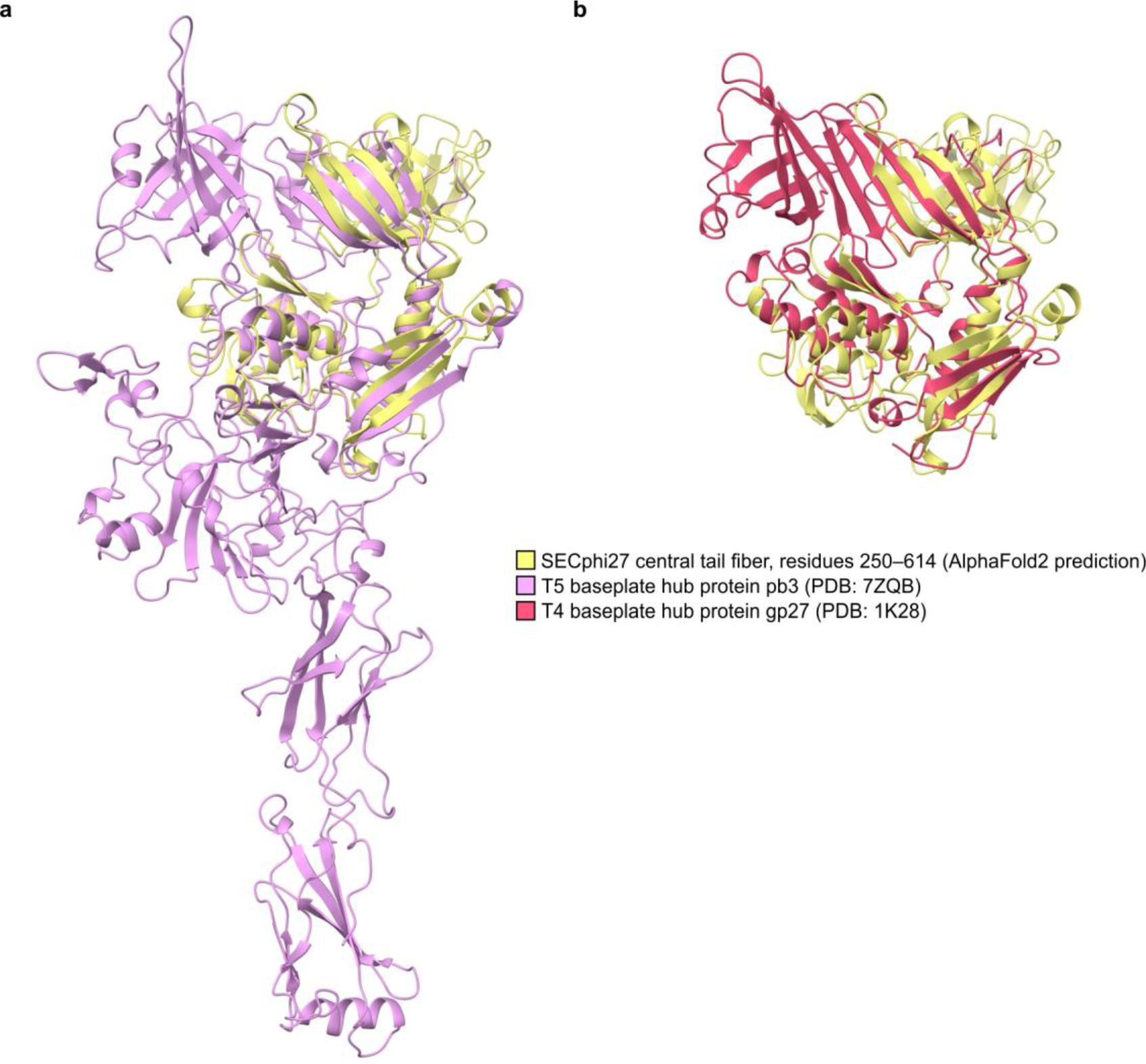
Tail structure proteins of different phages contain structurally similar domains. **a-b,** AlphaFold2^15^ prediction of residues 250–614 of the SECphi27 central tail fiber (related to Figure S5D) superimposed on (**a**) the cryo-EM structure of T5 pb3 (PDB: 7ZQB^31^; RMSD = 2.01 Å) or (**b**) the crystal structure of T4 gp27 (PDB: 1K28^51^; RMSD = 3.64 Å).

**Figure S7.**
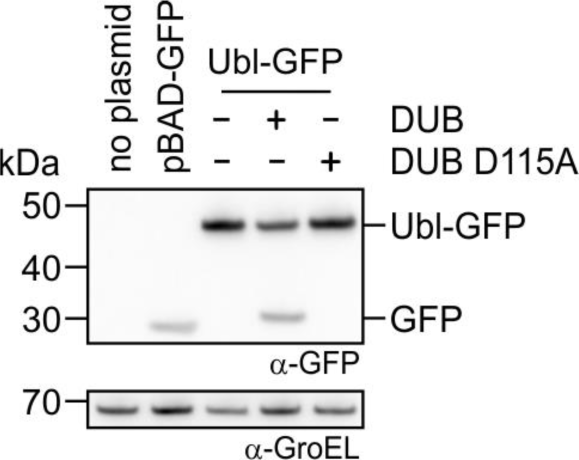
The DUB is able to cleave a Ubl fusion protein. Western blot of whole cell lysates of bacteria co-expressing a Ubl-GFP fusion protein and the DUB of the Bil system. GFP can be cleaved off the Ubl-GFP fusion protein by the DUB. GroEL was used as loading control.

